# Scaling law links plant growth variation to grain yield in wheat stands

**DOI:** 10.1101/2025.05.19.654792

**Authors:** Guy Golan, François Vasseur, Yongyu Huang, Kenan Tan, Victor O. Sadras, Cyrille Violle, Thorsten Schnurbusch

## Abstract

Growth rate, a fundamental biological trait influencing plant resource use, scales predictably with plant mass, following remarkably consistent allometric power laws shaped by biophysical constraints and natural selection. However, how these laws apply to crop plants shaped by artificial selection, and how they manifest in agronomic traits, remains undefined. Under controlled greenhouse conditions, we quantified the relationship between plant mass and growth rate in 195 European winter wheat cultivars. We uncovered genetic variation in allometry linked to plant size, where increased leaf allocation and faster development elevated allometric exponents. Phenotypic and genetic analyses revealed adaptive strategies, ranging from large, slow-growing genotypes that support reproductive initiation to small, fast-growing genotypes that enhance reproductive effort. A shared genetic basis—associated with *Photoperiod response-1* (*Ppd-1*)—linked growth allometry in the greenhouse to genotype-by-environment interactions for grain yield in field trials. This variation in growth allometry, shaped by breeding in diverse environments, reflects strategies that enhance adaptation to weather conditions. Our findings demonstrate that growth allometry is biologically robust and agronomically important because it scales to wheat yield under diverse, realistic field conditions.

## Introduction

Organisms acquire resources from their environment, process them through metabolism, and allocate them to various functions with implications for fitness. The rate of resource uptake and the expenditure of energy and resources are largely influenced by organism size (1, 2). Allometry describes the disproportionate change in an organism’s morphology, physiology, and life-history traits relative to changes in its size (3). Allometry can be examined across various contexts: during an organism’s growth and development (ontogenetic allometry), across individuals at the same developmental stage (static allometry), or across species (evolutionary allometry) (4). Allometric relationships often follow a power law, Y = αX^β^ where trait Y correlates with organism mass X, with α as a constant and β as the allometric exponent (3), typically linearized and analyzed on a log-log scale as log Y = log α+β log X.

Since Kleiber’s seminal work identified a ¾ allometric exponent between metabolic rate and body mass across animal species (5), the pervasive nature of quarter-power allometric scaling was observed across biological phenomena (6). Such scaling is mostly explained by the metabolic scaling theory (MST), which proposes that quarter-power allometric exponents result from an equilibrium between the scaling of hydraulic transport costs and surface area for resource exchange (1). In plants, substantial evidence supports relatively invariant MST predictions concerning metabolic and growth rate, resource allocation, and life history traits (7–12). Yet, other studies suggested that allometric exponents may vary among genotypes (11, 12), or deviate from MST predictions (13). Such deviations were suggested to stem from the intrinsic non-linearity of metabolic scaling (14), selection of different adaptive strategies (11, 12), environmental and physiological effects (15, 16), or analysis focused on very small plants (17, 18).

While MST predictions have been tested in wild plants and variations in allometric relationships have been explored, their validity and the implications of such variations for crop yield remain largely unknown. In nature, selection favors competitive phenotypes, whereas agronomic selection favors “communal” phenotypes that return higher yield per unit of land (19–21). This shift in selective pressure has driven allometric scaling divergences between wild plants and crop plants; for example, Green Revolution cereals have a higher allocation to reproduction at the expense of stems that increases yield per unit land area and reduces competitive ability for light (22). Considering the evidence that variation in metabolic scaling can evolve through natural selection or genetic drift (16), it is critical to know whether breeders have utilized such variation to increase yield or environmental adaptation. Moreover, identifying links between metabolic scaling and yield may inform crop improvement through targeted breeding of allometric traits (23).

MST predicts that plant growth rate (biomass produced per time unit) scales to the ¾ power of plant mass(10), illustrating that plants grow relatively slower but become more resource-efficient as they grow. In wild plants, such as *Arabidopsis thaliana,* genetic variation in growth allometry underlies a tradeoff between reproductive yield and stress resistance, where deviations from the ¾ allometric exponent reduce seed production in favor of stress tolerance (11). Similarly, leaf area allometry in wild tomato species varies around the ¾ allometric exponent, mediating a tradeoff between fecundity and drought response (24). If crops have a similar link between different fitness components, selecting growth allometry traits could improve yield across environments.

In addition, genetic variation in growth allometry likely reflects differences in biomass allocation among plant organs (8, 22). Biomass allocation is a dynamic process, with annual plants beginning their life cycle by investing primarily in roots and leaves. As ontogeny proceeds, allocation shifts to supporting structures such as stems and ultimately to reproduction (25). These transitions are influenced by the developmental trajectory and plant size, constraining organ growth, and thus resource use and reproductive growth (18, 26, 27). In cereals, grain yield is associated with the total biomass produced per area and the proportion allocated to grain, expressed as the harvest index (28). However, while the harvest index provides insights into the ultimate outcome of biomass allocation, it is a static measure that overlooks the dynamic allocation processes occurring throughout ontogeny, particularly during reproductive development, where resource allocation is crucial for grain yield. Importantly, the harvest index does not capture size-dependent variation in allocation of plant resources (29). Inflorescence fertility in cereal crops is determined by the dynamics of floret initiation and degeneration (30). An extended early growth phase increases the number of spikelet and floret primordia, thereby enhancing yield. However, it may also lead to larger stems, intensifying floret degeneration due to tradeoffs in allocation between reproduction, stem growth, accumulation of reserve carbohydrates, and maintenance (27, 31, 32). Therefore, a deeper understanding of allometry among organs during key reproductive developmental stages is essential for improving grain yield.

Here, we quantified the allometric relationship between growth rate and plant mass during ontogeny of 195 European elite winter wheat (*Triticum aestivum* L.) cultivars grown under controlled conditions, with a focus on the critical phase when the number of fertile florets per spike is determined. Phenotypic analysis and genome wide associations (GWA) scans revealed variation in allometry across genotypes, shaped by differences in developmental rate and resource allocation patterns, both of which share a common genetic basis with allometry. Moreover, we identified genetic links between reproductive development and allometry, which scaled to grain yield in independent, agronomically realistic field trials in France and Germany. Overall, our study identifies growth allometry as a physiological bridge that genetically and developmentally connects life-history traits, reproductive strategies, and environmental adaptation.

## Results

### Size differences contribute to variation in growth allometry

We examined growth allometry during the ontogeny of 195 winter wheat cultivars by studying the relationship between plant biomass and absolute growth rate (biomass accumulated over thermal time), which is expected to be proportional to metabolic rate (10). We utilized phenotypic data recorded during the spike growth period (33) reported by Guo et al. (34–36). Growth analysis began at the terminal spikelet (TS) stage, which marks the cessation of spikelet primordia initiation and the onset of floret initiation. Development of floret primordia progresses through the white anther (WA) stage, reaching its maximum number by the green anther (GA) stage. Following this, floret degeneration initiates around the yellow anther (YA) stage and can persist through the tipping (TP) and heading (HD) stages, continuing up to anthesis (AN) (Fig. S1).

We fitted a linear regression across developmental stages and cultivars, determining the allometric exponent β as the slope of the allometric function y = α + βx, where x and y represent the logarithms of plant biomass and growth rate, respectively. The allometric exponent was only slightly higher than predicted by MST and observed across land plants (β=0.77, [CI=0.76,0.78]), suggesting that wheat breeding did not change growth scaling dramatically. However, a quadratic model (AIC = −2438) explained ontogenetic allometry better than linear regression (AIC = −2404) (Fig. 1, Table S1), highlighting that variation in allometry associates with plant size. We employed the quadratic model y=−7.06+1.22x−0.029x^2^ to determine the allometric exponent of each cultivar by estimating the first derivative (β=1.229−0.0582X). The allometric exponent from the quadratic function gradually and significantly decreased with ontogeny, from TS (β mean=0.8±0.02) to AN (β mean=0.70±0.02), and the cultivar-specific allometric exponents were used in subsequent analyses.

**Fig. 1.**
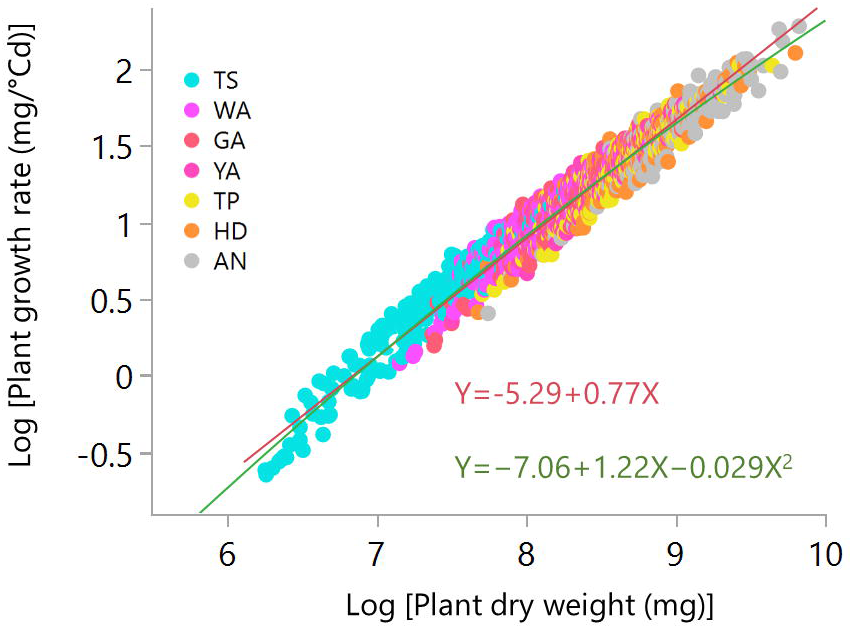
Variation in growth allometry in 195 hexaploid elite European winter wheat cultivars from Germany, France, Denmark and UK grown under controlled glasshouse conditions. The red line is the linear SMA regression and the green curve is the quadratic model of growth rate allometry across developmental stages. The data represent the means of the cultivars at each developmental stage: Terminal spikelet (TS), White anther (WA), Green anther (GA), Yellow anther (YA), Tipping (TP), Heading (HD), and Anthesis (AN).

### Variation in growth allometry is associated with resource use and reproductive strategies

We then explored how variation in growth allometry relates to wheat growth and resource use strategies throughout ontogeny. To do so, we examined its association with (i) growth duration (measured as the thermal time required to reach specific developmental stages), (ii) the cultivar’s relative growth rate, and (iii) biomass allocation to organs. Shorter growth duration was associated with a high allometric exponent (Fig. 2a). Moreover, the residuals from the quadratic function correlated negatively with thermal time, suggesting that the developmental rate exerts an additional influence on growth rate beyond the expected effects through plant size (Figs. S2a, S2b). The allometric exponent was also positively correlated with the plant’s relative growth rate (Fig S3). Overall, cultivars with a high allometric exponent prioritize rapid development, allowing plants to maximize their growth during short periods. Cultivars with low allometric exponents were associated with slow development, potentially more resource efficient and ensuing in a higher maximal size.

**Fig. 2.**
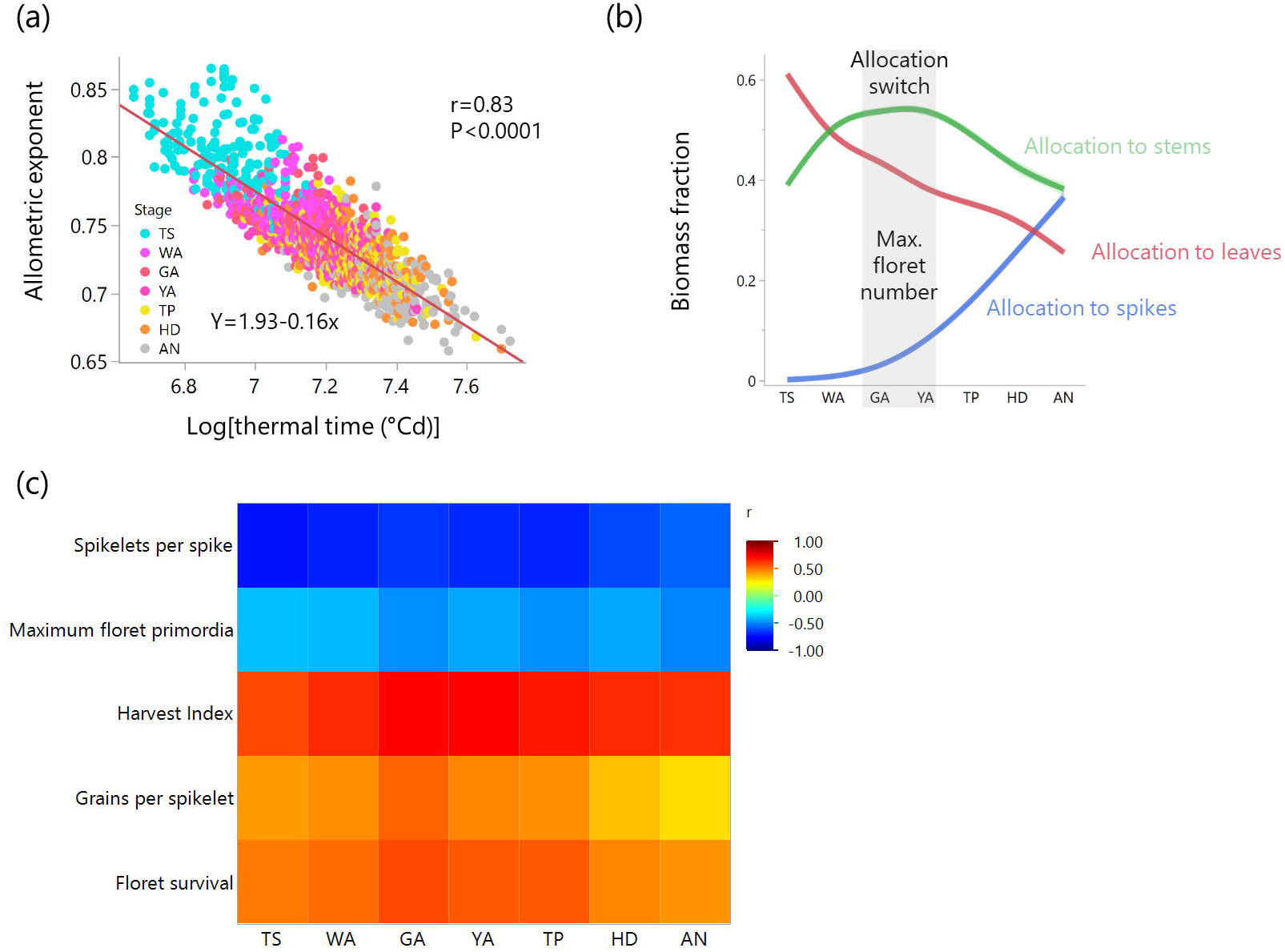
Growth allometry is associated with growth duration and reproductive traits. (a) Correlation between the allometric exponent and thermal time. (b) Biomass allocation patterns, modeled using a spline function with a 95% confidence interval, capturing nonlinear trends during ontogeny. (c) Correlations of reproductive traits with the cultivar’s growth rate allometric exponent at seven developmental stages: Terminal spikelet (TS), White anther (WA), Green anther (GA), Yellow anther (YA), Tipping (TP), Heading (HD), and Anthesis (AN). Data from195 hexaploid elite European winter wheat cultivars grown under controlled glasshouse conditions.

Biomass allocation to the leaves decreased during ontogeny (Fig. 2b) and became closely associated with the allometric exponent, starting from the WA stage, when substantial stem and spike growth commenced (Fig. S4). Allocation to the stem increased during the floret initiation period and peaked at the GA stage when number of primordia per spikelet peaked (33). Then, allocation shifted toward reproduction, and stem growth leveled off (Fig. 2b). The switch at the maximum floret primordia stage likely reflects a critical developmental checkpoint relevant to reproductive potential in wheat. Allocation to the stem before the maximum floret primordia stage promotes structural support and vascular capacity for transport, with labile carbohydrates contributing up to 40% of the total stem mass (37). Once the number of floret primordia is set, further growth shifts to ensure floret survival.

Allometric analysis of biomass allocation revealed that large cultivars allocate more biomass to the stems (inter-genotype allometric exponents >1) and less to the leaves and spikes (inter-genotype allometric exponents <1) (Table S2). Accordingly, small cultivars with high reproductive allocation exhibited higher floret survival rates and a higher harvest index, whereas large cultivars, with longer phases of reproductive development facilitated greater spikelet and floret initiation, increasing their potential grain number (Fig. S5).

Although growth allometry was associated with reproductive development across stages, the strength of these correlations varied (Fig. 2c), indicating that the effect of plant size on reproductive traits in wheat depends on the developmental stage. Overall, our results suggest that variation in growth allometry may reflect selection for resource use and reproductive strategies, and highlight the importance of plant size in determining yield at specific developmental windows.

### Genetic variation in growth allometry is associated with genotype-environment interactions for yield in independent field trials

Having found close relationships between reproductive traits and growth allometry, we examined whether growth allometry scales to grain yield under agronomically realistic conditions. This is biologically important and relevant in the context of plant breeding where phenotyping in simplified platforms, such as greenhouse or growth chambers, usually returns traits that do not correlate with grain yield in the field (38). To address this, we investigated grain yield data of the same wheat cultivars grown in eight distinct field environments, resulting from the combination of seasons and sites in Germany and France (39). Averaged yield varied from 79 dt/ha to 110 dt/ha across environments and from 85 dt/ha to 106 dt/ha across cultivars. The cultivar’s average grain yield across environments (BLUE) was unrelated to the allometric exponent and did show no trend with the cultivar’s year of registration (Figs. S6a, S6b). However, growth allometry was related to the country of origin of the cultivar. Analysis of variance indicated that the country of origin of the cultivar accounted for up to 23% (*P*<0.0001) of the variation in the allometric exponents (Table S3). French cultivars have allometric exponents significantly higher than Danish and German cultivars (Fig. 3a), suggesting that breeding for grain yield selected for environment-specific growth allometry, and highlighting the adaptive value of this trait.

**Fig. 3.**
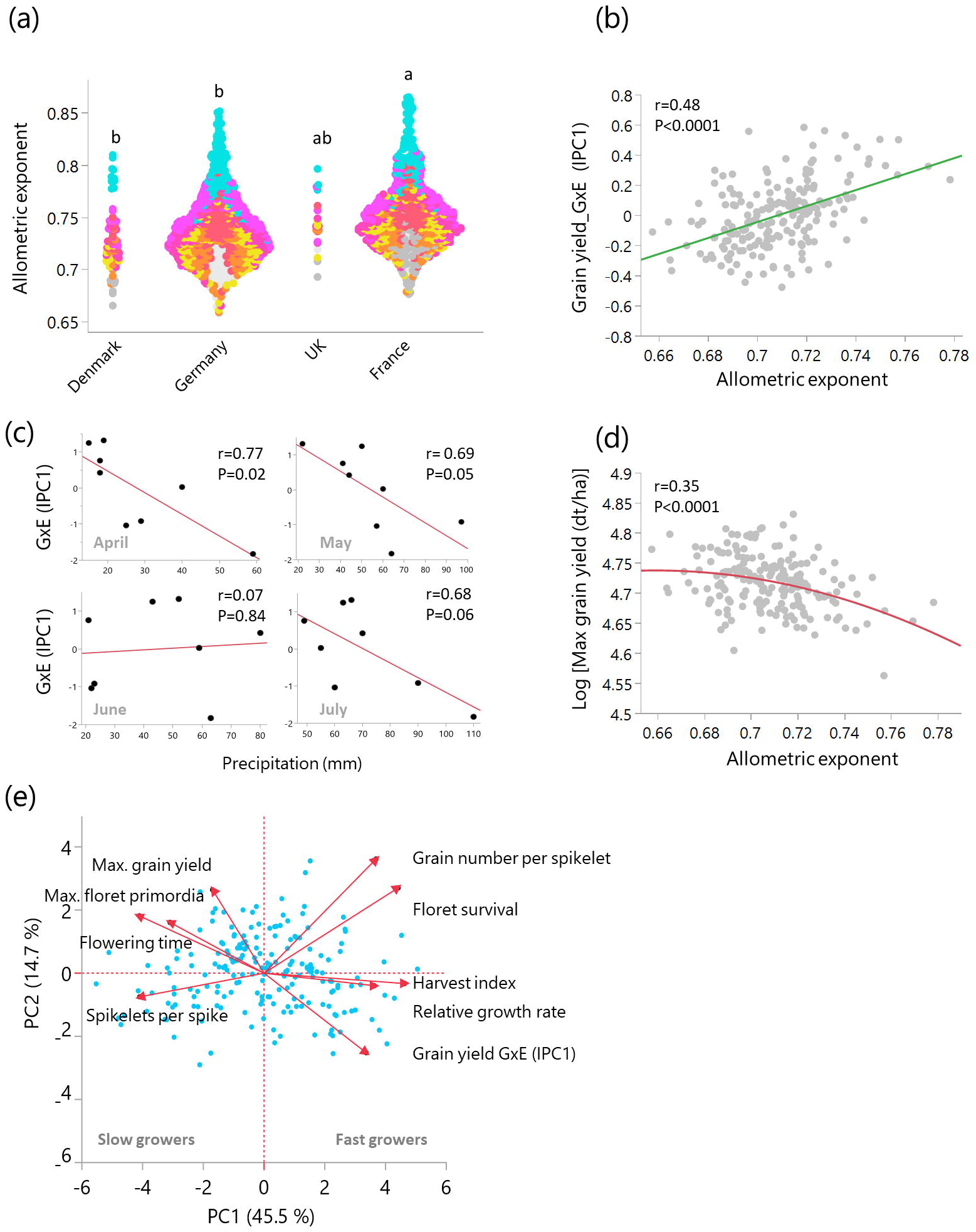
Growth rate allometry is associated with environmental adaptation in 195 hexaploid elite European winter wheat cultivars from Germany, France, Denmark and UK. (a) Associations between the cultivar’s country of registration and the allometric exponent at different developmental stages. Letters indicate statistically significant differences as determined by Tukey- Kramer HSD (P<0.05). (b) Correlation between the allometric exponent at anthesis in glasshouse- grown plants and GxE interactions for yield in field trials in Germany and France. (c) Correlation between the environment IPC1 and the monthly precipitation levels at the field sites. (d) Correlation between the allometric exponent at anthesis and the highest grain yield recorded across eight different environments. (e) PCA illustrating the differentiation of adaptive strategies shaped by growth allometry within the phenotypic space.

Thus, we tested whether the variation in growth allometry is related to the cultivar’s performance in contrasting environments. To address this, we used the AMMI (Additive Main Effects and Multiplicative Interaction) model (40) to dissect variation in genotype-environment interactions (GxE) into principal components for both genotypes and environments. This enabled us to investigate the associations between GxE interactions and the cultivar’s allometric exponent, as well as between GxE and weather conditions during the field trials. Interaction Principal Component 1 (IPC1) explained 31.7% of the GxE interactions and IPC2 accounted for 19.4% of the variation (Fig. S7). The allometric exponent recorded under controlled conditions explained 23% of the variation of the genotype IPC1 from independent field trials (Fig. 3b). Genotypes with higher allometric exponents performed relatively better in certain environments (2010JAN, 2009WOH), positioned farther from the origin in the AMMI biplot, indicating stronger genotype-environment interactions.

We then examined the relationship between the environmental IPC, which reflects the interactive forces of the environment, and the monthly rainfall and average temperatures at the trial sites. Environmental IPC1 scores were unrelated to monthly average temperatures (Fig. S8) and correlated with precipitation (Fig. 3c), highlighting the role of precipitation in shaping GxE interactions of rainfed crops. Considering also genotypes IPC1 correlations with the allometric exponents (Fig. 3b), we conclude that cultivars with high allometric exponents are better adapted to environments with lower rainfall during stem elongation (April-May) and grain filling (July). Taken together, genotype-dependent growth allometry plays an important role in environment-specific cultivar performance in major European wheat producing countries.

Notably, the allometric exponent correlated with the cultivar’s maximal grain yield recorded across environments, where lower allometric exponents, associated with a high number of spikelets and floret primordia, showed higher maximal yield. The observed relationship indicates that growth allometry impacts actual yield (Fig. 3d). Overall, the allometric exponent differentiated cultivars along a tradeoff of reproductive strategies. On the one side, fast-growing, smaller genotypes produce fewer spikelets per spike but maintain a higher proportion of viable florets per spikelet, return a higher harvest index, and yield well under low rainfall. On the other side, larger cultivars that prioritize initiation of spikelet and floret primordia achieve a higher maximum yield despite potentially lower floret survival (Fig. 3e).

### The genetic and molecular basis of growth allometry and its link to GxE interactions for grain yield in the field

To gain deeper insights into the genetics of the plant- and stand-level traits investigated, we conducted a genome-wide association scan with a focus on genomic regions associated with allometry, organ allocation patterns, growth rate, and growth duration measured in the greenhouse, and the GxE interactions for yield quantified in the field. We utilized the publicly available genotypic information of the cultivars (39) to analyze the population structure. Principal component analysis showed that PC1 explained 8.8% of the genetic variation and PC2 accounted for 4.3%. Although no clear distinction of subpopulations was observed within the panel, German cultivars displayed lower average PC1 values compared to French and Danish cultivars, suggesting genetic differentiation by country of registration (Figs. S9a, S9b). The allometric exponent was weakly correlated with PC1, PC3, and PC5 (Fig. S9c).

Our GWAS analysis identified 36 significant marker-trait associations (FDR<0.05) with growth allometry, corresponding to 19 QTL (Table S4). Of these, three QTL were also associated with organ allocation, growth rate, and duration, indicating a common genetic basis for growth allometry and resource-use strategies (Fig. 4a). One of the most prominent variants associated with growth allometry was a ∼2 kilo base pair (kb) deletion upstream of the *Photoperiod response 1* gene (*Ppd-D1*)(*41*) that was present in 15% of the tested cultivars, all of them registered in France. This variant confers photoperiod insensitivity, resulting in a higher developmental rate and shorter growth periods. Further GWAS and pairwise comparisons of different *Ppd-D1* alleles showed that this allelic variation influences key reproductive traits, such as spikelet number and harvest index, and is associated with relative growth rate, maximum floret primordia and floret survival (Table S4, Figs. 4b, 4c). The photoperiod-sensitive allele promoted spikelet and floret initiation, while the insensitive allele increased floret survival and harvest index at maturity.

**Fig. 4.**
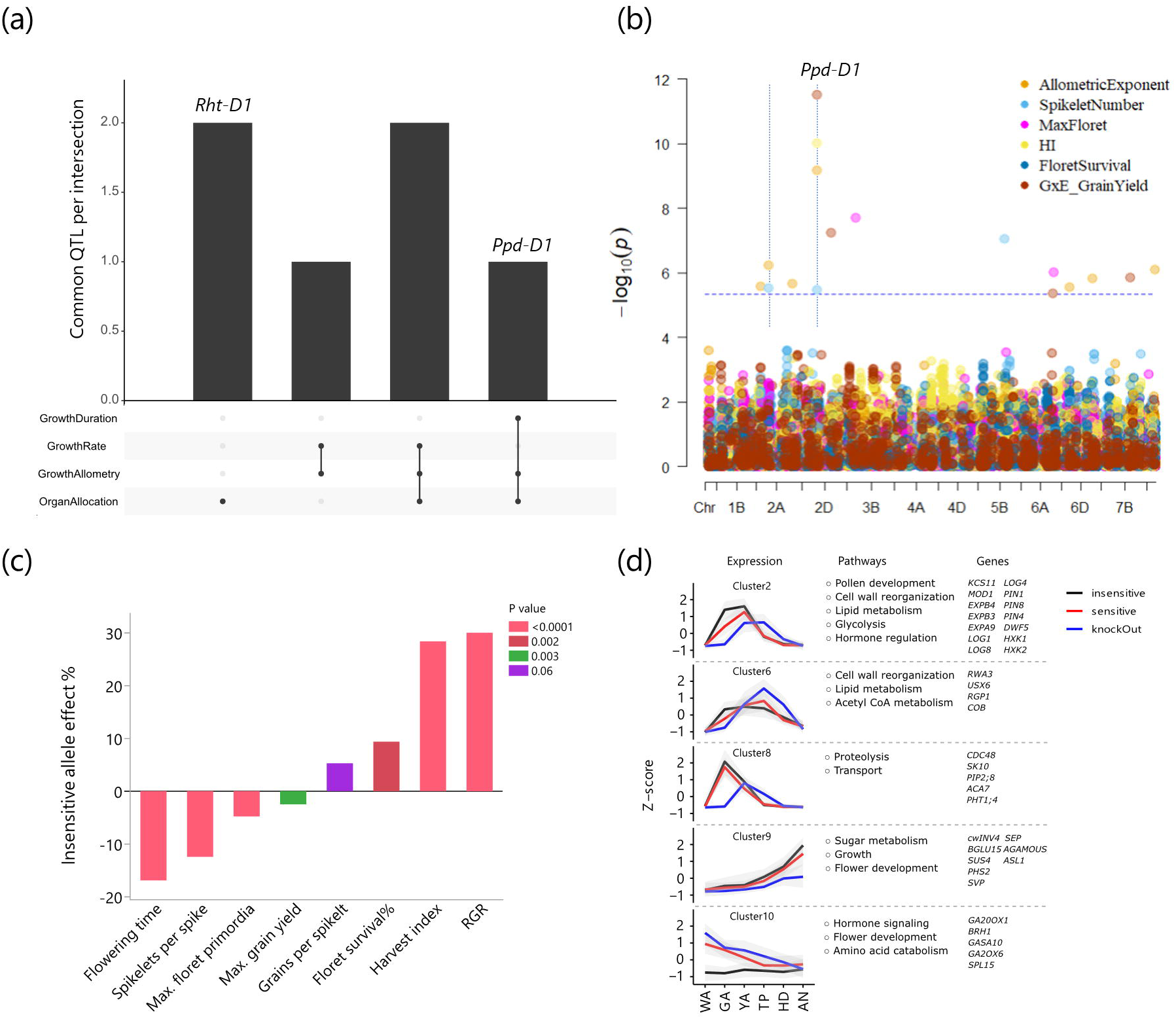
Genetic and molecular basis of growth rate allometry in 195 hexaploid elite European winter wheat cultivars. (a) UpSet plot depicting common QTL for allometry with growth traits. (b) Multi- trait Manhattan plot demonstrating shared genetic basis for growth allometry, reproductive traits, and grain yield GxE in eight environments. Horizontal dashed line indicates the Bonferroni correction threshold. Vertical dashed lines indicate common marker-trait associations. (c) The effects of the *Ppd-D1* insensitive allele on reproductive traits and relative growth rate (RGR). (d) *Ppd-1* allele driven spikelet gene expression dynamics from WA to AN. Gene clusters along with representative pathways and genes from enrichment analysis.

A previous study of *Ppd-D1* effects on reproductive development using near-isogenic lines and a loss of function mutant corroborated gene effects on spikelet number and fertility (42). Notably, *Ppd-D1*, which regulated resource allocation, growth duration, and reproductive strategies, also associated with GxE interactions for grain yield captured in principal components (IPC1) derived from the AMMI model (Fig. 4b, Table S4). This provides a genetic substrate to the phenotypic association between allometry traits and yield in the field.

To investigate how scaling of metabolism and growth are reflected at the molecular level and explore links with reproductive growth, we reanalyzed the spikelet transcriptome of three Paragon- derived genotypes with differing *Ppd-1* alleles across six developmental stages from WA to AN, as previously described (42). Pairwise comparisons among genotypes identified ∼19,000 differentially expressed genes (DEGs), grouped into ten clusters using the K-Medoids algorithm (Fig. S10, Table S5). Five clusters exhibited distinct allele effects on gene expression timing and intensity, driven primarily by heterochronic shifts (Fig. 4d). For example, cluster C2 showed early and higher expression in the photoperiod-insensitive genotype, peaking at GA, at the peak of floret primordia formation, and declining after YA. This cluster was enriched in genes related to cell wall modification (e.g., expansins), glycolysis (e.g., HXK1, HXK2), lipid metabolism (e.g., MOD1, KCS11), and cytokinin and brassinosteroid biosynthesis (e.g., LOGs, DWF5), reflecting increased metabolic investments that accelerate early spike growth. Similarly, cluster C6 displayed early induction of genes involved in cell wall organization, lipid biosynthesis, acetyl-CoA metabolism, and transmembrane transport in the insensitive genotype.

Cluster C8 showed specific expression of protein catabolism and membrane transport genes at GA in WT and NIL but delayed in the KO. Genes in cluster C9, associated with sugar metabolism (e.g., cwINV4, SUS4) and floral development (e.g., SEP3, SVP), exhibited the highest expression in the insensitive genotype and the lowest in KO. These patterns highlight the role of *Ppd-1* in metabolic scaling, enabling faster growth. In contrast, cluster C10 showed consistently low expression in the insensitive line, whereas WT and KO exhibited declining expression patterns. This cluster was enriched with gibberellin-related genes, potentially influenced by the high expression of cytokinin biosynthesis genes in the insensitive line during early spike growth. Considering the spikelet transcriptome alongside the performance of *Ppd-D1* across environments, we propose that the early metabolic upregulation observed in *Ppd-D1* insensitive genotypes represents an adaptive strategy for efficient resource utilization in environments where rapid reproductive development is adaptive, particularly under water deficit.

## Discussion

The metabolic and growth rates of plants influence their resource requirements (2). Understanding how growth rate changes throughout plant development is crucial for determining how resource use relates to growth patterns, reproductive development and performance in different environments, enabling more effective management strategies and predictive modeling (23, 43, 44).

Our large-scale dataset, encompassing high-resolution ontogenetic dynamics paired with field trial outcomes across eight contrasting environments, revealed developmental, physiological, genetic, and molecular connections between individual plant growth allometry under controlled conditions and grain yield in the field. By integrating ecological theory with crop science, we exemplify how artificial selection has, on the one hand, inadvertently shaped growth and metabolic scaling, advancing our understanding of allometry. On the other hand, our findings highlight the feasibility of applying allometric laws to better understand and improve crop performance.

The non-linear relationship between size and growth rate reflects variation in short-term growth dynamics among cultivars, variation that is averaged out in a linear regression across a 35-fold range of plant mass (Fig. 1). While likely not a direct target of selection, this variation is highly relevant for crop adaptation. Moreover, the robustness of allometry, i.e. scaling from controlled conditions to the field, opens new opportunities for screening breeding material for allometric traits at the individual-plant level to enhance genetic gain at the stand level (20, 38). Indeed, we found that breeders have leveraged genetic variation in shoot growth allometry to enhance grain yield in dry regions by altering developmental rates and organ allocation, thereby promoting yield components, such as floret survival and harvest index, while compromising on potential grain number.

Even though crop improvement can be considered as an empirical breeding and selection process, we did not observe time trends during the contemporary breeding of European cultivars (Fig. S6). This may be because variation in growth allometry may result from grain yield selections in specific environments. In France, larger latitudinal variation in climate and soil likely requires more environment-specific breeding. Small, fast-developing plants with high allometric exponents are agronomically adapted to the southern regions, where extended dry periods are common. On the other hand, large genotypes can return higher yield in wetter, higher-yielding environments of northern Europe. Long-term genetic yield improvements in Europe, the United States, and Argentina have been attributed to a balanced contribution of increased yield potential and maintenance breeding, ensuring cultivars remain adapted to evolving abiotic and biotic conditions (45), highlighting again a so far untapped potential for growth allometry in wheat breeding. Interestingly, a previous study using the complete GABI wheat panel investigated temperature responses related to stem elongation and found that French cultivars exhibit a more pronounced increase in elongation rate with rising temperatures than German cultivars (46). This response likely contributes to their fast growth, which is reflected in higher allometric exponents and adaptation to low rainfall shown in our study.

Allometry can be considered a form of phenotypic integration, arising from variations in developmental processes that lead to the covariation of multiple traits across organization levels (47, 48). Most tellingly, our genetic and phenotypic analysis identifies *Ppd-1* as a key regulator of vegetative and reproductive trait covariation integrated within growth allometry. *Ppd-1* functions within the photoperiod pathway, regulating genes underlying developmental timing (35, 49), thereby determining developmental windows for organ growth. Moreover, *Ppd-1* directly activates the gibberellin biosynthesis pathway in internodes, thereby influencing elongation processes (31). Consequently, as shown here, allelic *cis*-variation at *Ppd-1* associates with variation in resource allocation and growth rate, indicating that *Ppd-1* is thus a major locus for growth allometry in wheat.

Previous studies found that gradual increases of flowering signals during reproductive development modulates fertility and grain yield (49–51). The photoperiod-sensitive alleles (lower signal intensity) allow longer periods of spikelet initiation, whereas the stronger flowering signal in the photoperiod-insensitive lines elicits the early induction of metabolic and floral development genes that promote floret survival (Figs. 4d). In line with our findings, *Ppd-1* plays a role in stabilizing barley (*Hordeum vulgare* L.) reproductive output under high temperatures by boosting metabolic genes in the spike, allowing acceleration of reproductive development to mitigate temperature effects on floret fertility (52).

Growth allometry is a key integrator not only for developmental but also for environmental signals. Environmental factors significantly affect plant size and consequently affect resource allocation in the plant (53). Here, the interaction between plant size and the developmental stage plays an important role. For example, a small plant before the allocation switch (Fig. 2b) will reduce stem allocation and increase the proportion of leaves. Beyond the allocation switch, size reductions will affect reproductive allocation, and the proportion of vegetative parts will become larger. However, the genotype’s allometric trajectory for reproductive allocation during this critical growth phase influences the degree of phenotypic plasticity in reproductive growth. A steeper increase in reproductive vs vegetative growth will promote yield but will also make the genotype more sensitive to environmental stresses (29, 44, 54).

Overall, our study bridges a gap between fundamental understanding of plant allometry and agronomic applications. We advocate for the inclusion of growth allometry in crop phenotyping within a breeding context(23), since we showcase that it scales to grain yield under relevant field conditions, and is further supported by genetic and metabolic evidence. Combining genetic information on allometric traits with environmental, climate, soil, and management data enables better predictions, modeling, and selection of high-yielding genotypes across environments.

## Methods

### Growth conditions and phenotypic measurements

Phenotypic data for growth analysis were collected from 195 hexaploid elite winter wheat cultivars, a subset of the GABI wheat panel (39), grown under controlled greenhouse conditions at IPK Gatersleben, Germany, as previously described(34–36). The investigated panel included cultivars from Germany (n=93), France (n=87), Denmark (n=10), and the UK (n=3), and one cultivar each from the Czech Republic and Austria. Seedlings were vernalized for 63 days at 4°C and then cultivated under long days with a 16:8 h photoperiod and day/night temperatures of 20°C and 6°C, respectively. Starting two weeks after vernalization, plants were dissected every two days to determine the development stage using the Kirby scale (55). The developmental stages of the cultivar and the thermal time to reach the stages were assessed using at least three randomly selected plants. A cultivar was considered to be at the TP, HD, or AN stage when at least half of the plants reached the corresponding stage. Once the plants reached a specific stage, three plants of each cultivar were randomly selected and separated for tillers, and the main culm and its biomass were determined. The main culm was used to determine organ weight, spikelet number, maximum floret primordia number, and the percentage of floret survival (grains per spikelet/ maximum floret primordia). Thermal time (°Cd) was used to quantify growth duration to reach each stage. It was calculated as the sum of cumulative daily thermal, calculated as T_average_ - T_base_. The base temperature (T_base_) was assumed to be 0°C and average temperatures (T_average_) were calculated as [(T_max_ + T_min_)/2].

### Growth analysis

The absolute growth rate for each cultivar at each developmental stage was calculated as the total plant biomass accumulated divided by thermal time of the growth period. Standard Major Axis (SMA) regressions were conducted using the ‘SMATR’ package(56) in R. The relative growth rate (RGR) for each cultivar was extracted from a two-parameter exponential growth model implemented in JMP17 (SAS Institute, Cary, NC). The model is defined as:

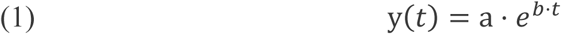

where a represents the plant’s initial biomass at TS, b is the estimated relative growth rate, t is thermal time, and e is the base of natural logarithms.

### Grain yield analysis

Grain yield data for the same cultivars examined in the greenhouse were collected from plots (5- 6.75m^2^) in eight field experiments conducted in Germany and France during 2009–2010, as previously described (39). Environment-specific and average yield (BLUE) were obtained from the e!DAL-PGP-Repository (https://doi.org/10.5447/ipk/2022/18). Genotype-by-environment (GxE) interactions for grain yield were analyzed using the Additive Main Effects and Multiplicative Interaction (AMMI) model, implemented in Genstat (VSN International, Hemel Hempstead, UK), following the approach described by Malosetti et al. (57). The AMMI model first applies analysis of variance (ANOVA) to partition the total variance into genotypic, environmental, and GxE interaction components. Then, Principal Component Analysis (PCA) is applied to the GxE interaction term, extracting Interaction Principal Components (IPCs) that summarize the interaction structure. The model is expressed as:

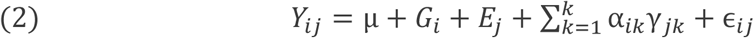

where *Y*_*ij*_ represents the observed response of genotype *i* in environment *j*, *G_i_* and *E_j_* are the additive main effects of genotype and environment, respectively, and the interaction term is decomposed into K significant Interaction Principal Components (IPC). Each IPC consists of a genotypic sensitivity score α_*ik*_ and an environmental score γ_*jk*_. The IPC scores for genotypes and environments were visualized using biplots and utilized for subsequent analyses. Environmental data of temperature and precipitation at trial sites were obtained from a previous study (58).

### Association mapping

Genotypic data from the 35K and 90K iSELECT SNP arrays, along with marker oligo sequences were obtained from the Dryad repository (39). Markers were mapped to the Chinese Spring reference genome (version 1.1) using BLASTN, retaining only the genomic positions with the smallest E-values. Markers that could not be unambiguously positioned on the genome were excluded from the GWAS analysis. Genotypic data for functional gene markers at the *Ppd-D1, Rht-B1, Rht-D1*, and *TaGW2* loci, previously reported (59), were also included in the analysis. After filtering out SNPs with a minor allele frequency (MAF) <0.05, a total of 10,781 biallelic markers were used for the GWAS.

Marker-trait associations were analyzed using the R package GAPIT3(60)(V3.4.0) with two multi- locus models: FarmCPU (61) and BLINK (62). Population structure was assessed using principal component analysis (PC1–PC5), of the genotypic data. Of these, the first three principal components were used as covariate fixed effects in the GWAS models to account for population structure. Markers with an FDR-adjusted (B&H) (63) P-value <0.05 were considered statistically significant.

### Transcriptome analysis

The RNA sequencing was done with five biological replicates where each sample consists of spikelets from at least three plants. Sample collection, and preparation were mentioned previously (42). The sequencing raw reads were first subjected to adapter trimming and quality control filtering using Fastp (64) (v.0.20.0) with standard parameters and workflows. We used Chinese Spring gene annotation V1.1 as a reference to estimate read abundance using the software Kallisto(65). Gene-level abundance (transcripts per million, TPM) and counts were summarized with tximport (66) (v3.14) DESeq2 tool (67) was used for differentially expressed genes (DEG) analysis using genes with read counts ≥10 in at least 3 samples. A pairwise comparison was used to determine DEG by considering genotypic effect across stages (18 comparisons). Genes with |log2 FC| ≥ 1 and a Benjamini-Hochberg FDR-adjusted P-value < 0.05 between genotypes were considered as DEGs, resulting in total 19,022 dynamically expressed genes (DYGs) (Table S5). Averaged TPMs from the DYGs were Z-scored and clustered using the K-medoids method with the PAM algorithm implemented in the R package cluster (68) (2.1.7). The number of partitions to be clustered was set at 10 clusters based on Gap-Statistics. Euclidean distance was used for the clustering. Enrichment analysis of GO terms (biological processes) was conducted using the closest *Arabidopsis thaliana* homologs. The TGT database (69) was employed to identify one reciprocal best hit (RBH) *Arabidopsis* homolog per wheat gene ID. GO term enrichment analysis was performed and summarized using Metascape (70) (http://metascape.org) with default parameters.

## Acknowledgements

The authors gratefully acknowledge Dr Zifeng Guo, now at The Key Laboratory of Plant Molecular Physiology, Chinese Academy of Sciences (CAS), Beijing, China, and Mechthild Pürschel for conducting the controlled greenhouse experiments and collecting the phenotypic data analyzed in this study. We thank Hannah Schneider for valuable comments on an earlier version of the manuscript.

## Author Contributions

GG: Conceptualization, data analysis and writing of the first draft. FV: Writing, manuscript review and editing. YH: Transcriptome analysis, writing, manuscript review and editing. KT: Bioinformatic processing of transcriptome data. VS: Writing, manuscript review and editing. CV: Writing, manuscript review and editing. TS: Funding acquisition, writing, manuscript review and editing.

---

**Fig. S1** Schematic representation of the initiation and degenerations of floret primordia along the spikelet development scale defined by Kirby and Appleyard (1981).

**Fig. S2** Residuals analysis of ontogenetic allometry. (a) Predicted growth rate vs residuals of the quadratic model presented in Fig. 1. (b) Correlations between growth duration and residuals. All correlation P-values were below 0.0001.

**Fig. S3** Growth allometry correlates with relative growth rate. (a) Correlations between the cultivar’s relative growth rate and the allometric exponent at different developmental stages.

**Fig. S4** Correlation between the leaf mass fraction (LMF) and the allometric exponent throughout ontogeny.

**Fig. S5** Development of reproductive traits and their associations with the growth rate allometric exponent during yield-defining stages. (a) Correlation between the allometric exponent at TS and the number of spikelets per spike. (b) Correlation between the allometric exponent at GA and the maximum number of floret primordia. (c) Correlation between the allometric exponent at GA and the percentage of floret primordia that ended up setting grains (d) Correlation between the allometric exponent at GA and the number of grains per spikelet. (e) Correlation between the allometric exponent at YA and the harvest index at maturity.

**Fig. S6** Growth allometry do not affect stable grain yield effects. (a) Relationships between the allometric exponent at different stages and the cultivar’s grain yield per area, averaged across eight environments. (b) Relationships between the cultivar’s year of release and the allometric exponent at different developmental stages.

**Fig. S7** Visualization of G × E interactions for grain yield across eight environments in Germany and France. A biplot using symmetric scaling represents cultivars as blue circles and environments as green diamonds. Cultivars near the origin are less sensitive to environmental interactions, whereas those located farther from the origin exhibit specific adaptation.

**Fig. S8** Correlations between the average temperature (Celsius degrees) in the eight field trials with the interaction principal components (IPC) extracted from the AMMI model.

**Fig. S9** Population structure of the GABI wheat panel and its association with the country of origin and the scaling exponent. (a) Biplot of the population structure of the GABI wheat panel. (b) Association between the cultivar’s country of registration and the population structure principal components. (c) Relationships between the scaling exponent at different stages and the population structure principal components.

**Fig. S10** Gene expression in the spikelets of near-isogenic lines with differing *Ppd-1* alleles, observed from WA to AN. (a) The number of up regulated and down regulated genes between genotypes at different developmental stages. (b) Clusters of DEGs expression during spikelet growth.

## References

1. G. B. West, J. H. Brown, B. J. Enquist, A general model for the origin of allometric scaling laws in biology. Science 276, 122–126 (1997).

2. J. H. Brown, J. F. Gillooly, A. P. Allen, V. M. Savage, G. B. West, Toward a metabolic theory of ecology. Ecology 85, 1771–1789 (2004).

3. J. S. Huxley, Constant Differential Growth-Ratios and their Significance. Nature 114, 895–896 (1924).

4. C. Pélabon et al., On the relationship between ontogenetic and static allometry. The American Naturalist 181, 195–212 (2013).

5. M. Kleiber, Body size and metabolism. Hilgardia 6, 315–353 (1932).

6. V. M. Savage et al., The predominance of quarter-power scaling in biology. Functional Ecology 18, 257–282 (2004).

7. B. J. Enquist, J. H. Brown, G. B. West, Allometric scaling of plant energetics and population density. Nature 395, 163–165 (1998).

8. B. J. Enquist, K. J. Niklas, Global allocation rules for patterns of biomass partitioning in seed plants. Science 295, 1517–1520 (2002).

9. N. Marbà, C. M. Duarte, S. Agustí, Allometric scaling of plant life history. Proceedings of the National Academy of Sciences 104, 15777–15780 (2007).

10. K. J. Niklas, B. J. Enquist, Invariant scaling relationships for interspecific plant biomass production rates and body size. Proceedings of the National Academy of Sciences 98, 2922–2927 (2001).

11. F. Vasseur et al., Adaptive diversification of growth allometry in the plant *Arabidopsis thaliana*. Proceedings of the National Academy of Sciences 115, 3416–3421 (2018).

12. F. Vasseur, C. Violle, B. J. Enquist, C. Granier, D. Vile, A common genetic basis to the origin of the leaf economics spectrum and metabolic scaling allometry. Ecology Letters 15, 1149–1157 (2012).

13. P. B. Reich, M. G. Tjoelker, J.-L. Machado, J. Oleksyn, Universal scaling of respiratory metabolism, size and nitrogen in plants. Nature 439, 457–461 (2006).

14. T. Kolokotrones, S. Van, E. J. Deeds, W. Fontana, Curvature in metabolic scaling. Nature 464, 753–756 (2010).

15. D. A. Coomes, E. R. Lines, R. B. Allen, Moving on from Metabolic Scaling Theory: hierarchical models of tree growth and asymmetric competition for light. Journal of Ecology 99, 748–756 (2011).

16. D. S. Glazier, Beyond the ‘3/4-power law’: variation in the intra-and interspecific scaling of metabolic rate in animals. Biological Reviews 80, 611–662 (2005).

17. B. J. Enquist et al., Does the exception prove the rule? Nature 445, E9–E10 (2007).

18. H. Poorter et al., How does biomass distribution change with size and differ among species? An analysis for 1200 plant species from five continents. New Phytologist 208, 736–749 (2015).

19. R. F. Denison, E. T. Kiers, S. A. West, Darwinian agriculture: when can humans find solutions beyond the reach of natural selection? Q Rev Biol 78, 145–168 (2003).

20. G. Golan, R. Abbai, T. Schnurbusch, Exploring the trade-off between individual fitness and community performance of wheat crops using simulated canopy shade. Plant, Cell & Environment 46, 3144–3157 (2023).

21. J. Weiner, Y.-L. Du, C. Zhang, X.-L. Qin, F.-M. Li, Evolutionary agroecology: individual fitness and population yield in wheat (*Triticum aestivum*). Ecology 98, 2261–2266 (2017).

22. G. Golan, J. Weiner, Y. Zhao, T. Schnurbusch, Agroecological genetics of biomass allocation in wheat uncovers genotype interactions with canopy shade and plant size. New Phytologist, 107–120 (2024).

23. A. J. Westgeest et al., An allometry perspective on crops. New Phytologist (2024).

24. C. D. Muir, M. Thomas-Huebner, Constraint around Quarter-Power Allometric Scaling in Wild Tomatoes (Solanum sect. Lycopersicon; Solanaceae). The American Naturalist 186, 421–433 (2015).

25. J. L. Harper, J. Ogden, The reproductive strategy of higher plants: I. The concept of strategy with special reference to Senecio vulgaris L. The Journal of Ecology, 681–698 (1970).

26. C. Körner, Some often overlooked plant characteristics as determinants of plant growth: a reconsideration. Functional ecology, 162–173 (1991).

27. C. Rivera-Amado et al., Optimizing dry-matter partitioning for increased spike growth, grain number and harvest index in spring wheat. Field Crops Research 240, 154–167 (2019).

28. C. Donald, J. Hamblin, The biological yield and harvest index of cereals as agronomic and plant breeding criteria. Advances in agronomy 28, 361–405 (1976).

29. X.-l. Qin, J. Weiner, L. Qi, Y.-c. Xiong, F.-m. Li, Allometric analysis of the effects of density on reproductive allocation and Harvest Index in 6 varieties of wheat (Triticum). Field Crops Research 144, 162–166 (2013).

30. Y. Huang, T. Schnurbusch, The birth and death of floral organs in cereal crops. Annual Review of Plant Biology 75, 427–458 (2024).

31. Y. Huang et al., Dynamic phytomeric growth contributes to local adaptation in barley. Molecular Biology and Evolution 41, msae011 (2024).

32. K. H. M. Siddique, E. J. M. Kirby, M. W. Perry, Ear: Stem ratio in old and modern wheat varieties; relationship with improvement in number of grains per ear and yield. Field Crops Research 21, 59–78 (1989).

33. Z. Guo, T. Schnurbusch, Variation of floret fertility in hexaploid wheat revealed by tiller removal. Journal of experimental botany 66, 5945–5958 (2015).

34. Z. Guo et al., Genome-wide association analyses of 54 traits identified multiple loci for the determination of floret fertility in wheat. New Phytologist 214, 257–270 (2017).

35. Z. Guo, D. Chen, M. S. Röder, M. W. Ganal, T. Schnurbusch, Genetic dissection of pre-anthesis sub-phase durations during the reproductive spike development of wheat. The Plant Journal 95, 909–918 (2018).

36. Z. Guo et al., Genome-wide association analyses of plant growth traits during the stem elongation phase in wheat. Plant Biotechnology Journal 16, 2042–2052 (2018).

37. V. O. Sadras, Evolutionary and ecological perspectives on the wheat phenotype. Proceedings of the Royal Society B 288, 20211259 (2021).

38. V. O. Sadras, Effective phenotyping applications require matching trait and platform and more attention to theory. Front Plant Sci 10, 1339 (2019).

39. A. Gogna, A. W. Schulthess, M. S. Röder, M. W. Ganal, J. C. Reif, Gabi wheat a panel of European elite lines as central stock for wheat genetic research. Scientific Data 9, 538 (2022).

40. R. W. Zobel, M. J. Wright, H. G. Gauch Jr, Statistical analysis of a yield trial. Agronomy journal 80, 388–393 (1988).

41. J. Beales, A. Turner, S. Griffiths, J. W. Snape, D. A. Laurie, A pseudo-response regulator is misexpressed in the photoperiod insensitive *Ppd-D1a* mutant of wheat (*Triticum aestivum L*.). Theoretical and Applied Genetics 115, 721–733 (2007).

42. Y. Liu et al., Ppd-1 remodels spike architecture by regulating floral development in wheat. BioRxiv, 2020.2005. 2011.087809 (2020).

43. G. Lemaire, T. Sinclair, V. Sadras, G. Bélanger, Allometric approach to crop nutrition and implications for crop diagnosis and phenotyping. A review. Agronomy for Sustainable Development 39, 27 (2019).

44. J. Weiner, Y.-L. Du, Y.-M. Zhao, F.-M. Li, Allometry and yield stability of cereals. Frontiers in Plant Science 12 (2021).

45. P. Grassini, et al., Maintenance breeding and breeding for yield potential equally contribute to genetic improvement in wheat yield. (2025).

46. L. Roth et al., High-throughput field phenotyping reveals that selection in breeding has affected the phenology and temperature response of wheat in the stem elongation phase. Journal of Experimental Botany 75, 2084–2099 (2023).

47. B. Hallgrímsson et al., Integration and the developmental genetics of allometry. Integrative and Comparative Biology 59, 1369–1381 (2019).

48. F. Vasseur, A. J. Westgeest, D. Vile, C. Violle, Solving the grand challenge of phenotypic integration: allometry across scales. Genetica 150, 161–169 (2022).

49. A. Gauley et al., Photoperiod-1 regulates the wheat inflorescence transcriptome to influence spikelet architecture and flowering time. Current Biology 34, 2330–2343. e2334 (2024).

50. L. M. Shaw et al., FLOWERING LOCUS T2 regulates spike development and fertility in temperate cereals. Journal of Experimental Botany 70, 193–204 (2018).

51. G. V. Yoshikawa, S. A. Boden, Finding the right balance: The enduring role of florigens during cereal inflorescence development and their influence on fertility. Current Opinion in Plant Biology 79, 102539 (2024).

52. T. Lan et al., PHOTOPERIOD 1 enhances stress resistance and energy metabolism to promote spike fertility in barley under high ambient temperatures. Plant Physiology 197 (2025).

53. J. Weiner, Allocation, plasticity and allometry in plants. Perspectives in Plant Ecology, Evolution and Systematics 6, 207–215 (2004).

54. Y.-L. Du et al., Yield components, reproductive allometry and the tradeoff between grain yield and yield stability in dryland spring wheat. Field Crops Research 257, 107930 (2020).

55. E. M. Kirby, M. Appleyard, Cereal development guide. (1984).

56. D. I. Warton, R. A. Duursma, D. S. Falster, S. Taskinen, smatr 3– an R package for estimation and inference about allometric lines. Methods in Ecology and Evolution 3, 257–259 (2012).

57. M. Malosetti, J.-M. Ribaut, F. A. van Eeuwijk, The statistical analysis of multi-environment data: modeling genotype-by-environment interaction and its genetic basis. Frontiers in Physiology 4 (2013).

58. C. D. Zanke et al., Whole genome association mapping of plant height in winter wheat *(Triticum aestivum* L.). PLOS ONE 9, e113287 (2014).

59. C. D. Zanke et al., Analysis of main effect QTL for thousand grain weight in European winter wheat (Triticum aestivum L.) by genome-wide association mapping. Front Plant Sci 6, 644 (2015).

60. J. Wang, Z. Zhang, GAPIT Version 3: Boosting power and accuracy for genomic association and prediction. Genomics Proteomics Bioinformatics 19, 629–640 (2021).

61. X. Liu, M. Huang, B. Fan, E. S. Buckler, Z. Zhang, Iterative usage of fixed and random effect models for powerful and efficient genome-wide association studies. PLOS Genetics 12, e1005767 (2016).

62. M. Huang, X. Liu, Y. Zhou, R. M. Summers, Z. Zhang, BLINK: a package for the next level of genome-wide association studies with both individuals and markers in the millions. GigaScience 8 (2018).

63. Y. Benjamini, Y. Hochberg, Controlling the false discovery rate: a practical and powerful approach to multiple testing. Journal of the Royal statistical society: series B (Methodological) 57, 289–300 (1995).

64. S. Chen, Y. Zhou, Y. Chen, J. Gu, fastp: an ultra-fast all-in-one FASTQ preprocessor. Bioinformatics 34, i884–i890 (2018).

65. N. L. Bray, H. Pimentel, P. Melsted, L. Pachter, Near-optimal probabilistic RNA-seq quantification. Nature biotechnology 34, 525–527 (2016).

66. C. Soneson, M. I. Love, M. D. Robinson, Differential analyses for RNA-seq: transcript-level estimates improve gene-level inferences. F1000Res 4, 1521 (2015).

67. M. I. Love, W. Huber, S. Anders, Moderated estimation of fold change and dispersion for RNA-seq data with DESeq2. Genome biology 15, 1–21 (2014).

68. M. Maechler, Cluster: cluster analysis basics and extensions. R package version 2.0. 7–1 (2018).

69. Y. Chen et al., A collinearity-incorporating homology inference strategy for connecting emerging assemblies in the triticeae tribe as a pilot practice in the plant pangenomic era. Molecular Plant 13, 1694–1708 (2020).

70. Y. Zhou et al., Metascape provides a biologist-oriented resource for the analysis of systems-level datasets. Nature communications 10, 1523 (2019).

