## Supplementary figures for "Scaling law links plant growth variation to grain yield in wheat stands"

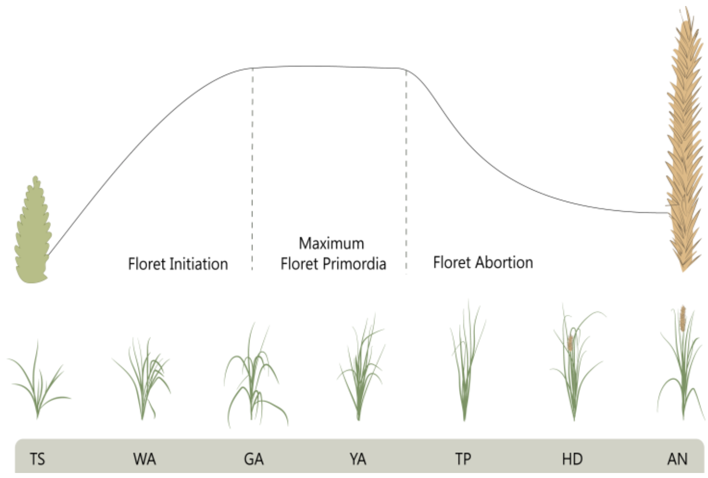


**Fig. S1** Schematic representation of the initiation and degenerations of floret primordia along the spikelet development scale defined by Kirby and Appleyard (1984).

(a)

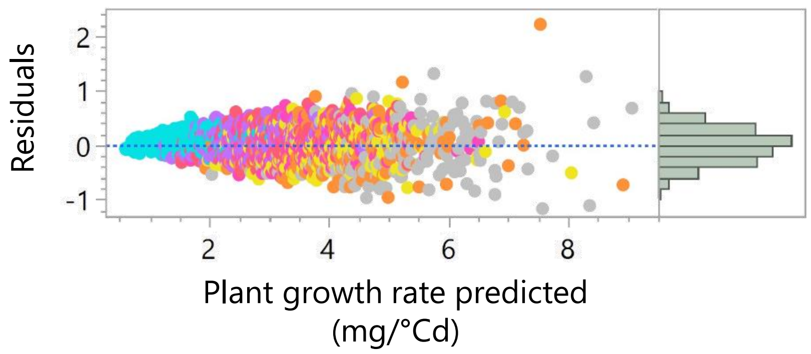


(b)

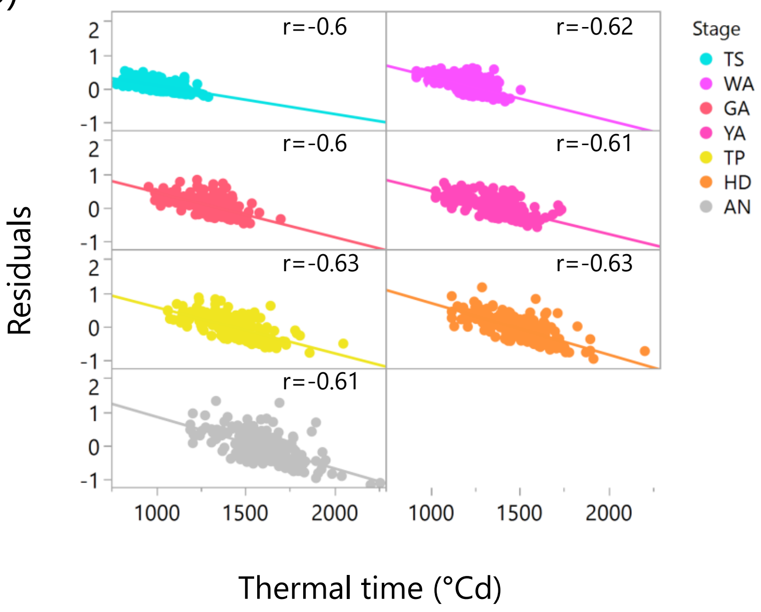


**Fig. S2** Residuals analysis of ontogenetic allometry. (a) Predicted growth rate vs residuals of the quadratic model presented in Fig. 1. (b) Correlations between growth duration and residuals. All correlation P-values were below 0.0001.

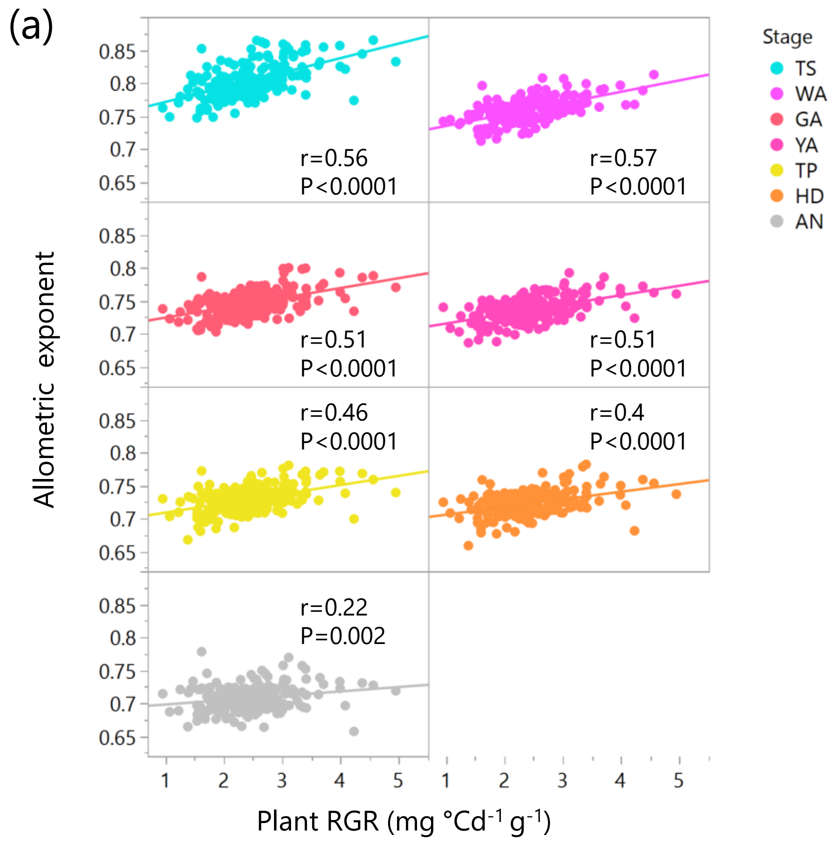


**Fig. S3** Growth allometry correlates with relative growth rate. (a) Correlations between the cultivar's relative growth rate and the allometric exponent at different developmental stages.

(a)

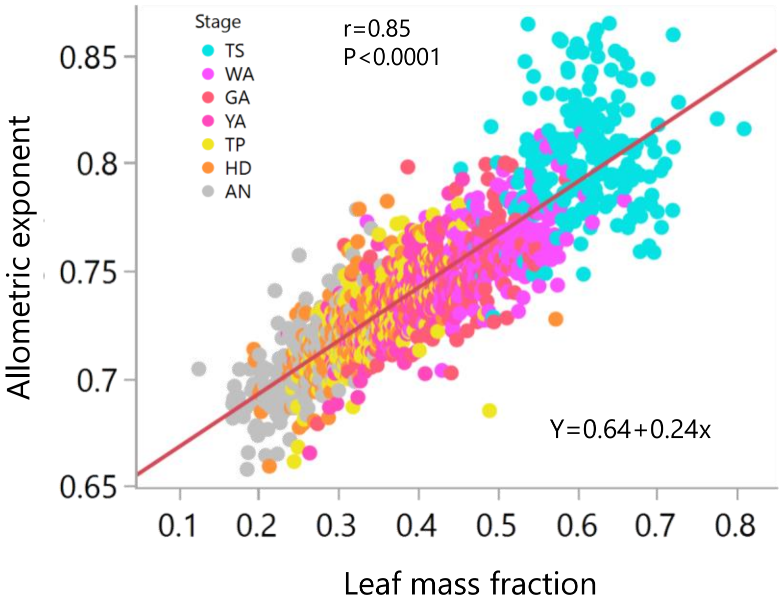


**Fig. S4** Correlation between the leaf mass fraction (LMF) and the allometric exponent throughout ontogeny.

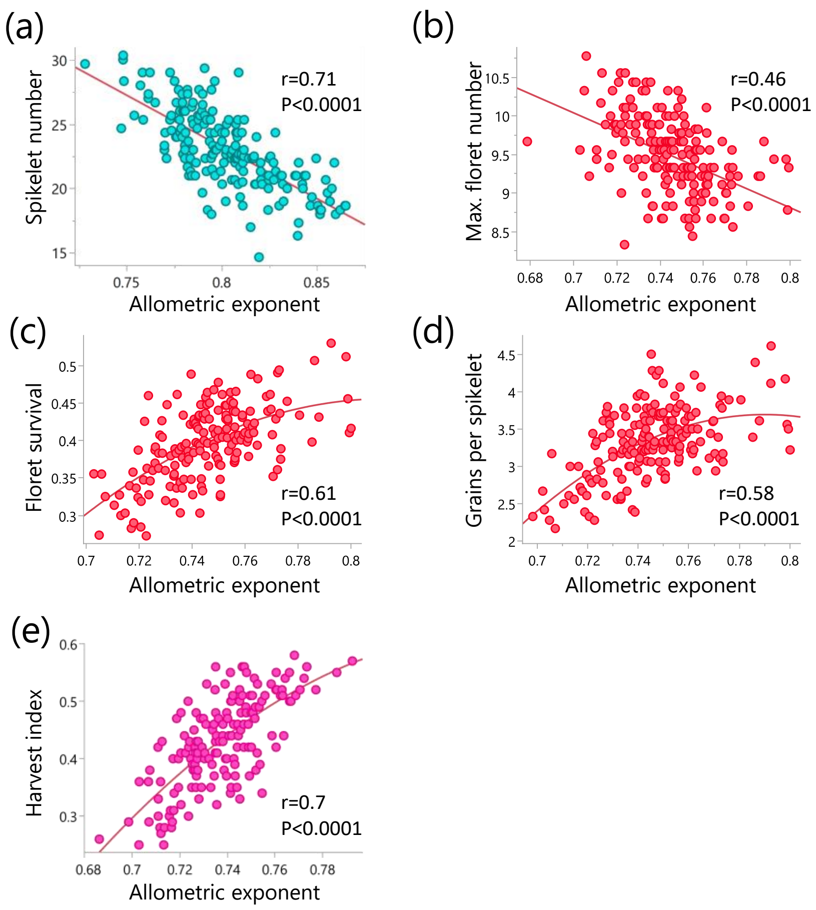


**Fig. S5** Development of reproductive traits and their associations with the growth rate allometric exponent. (a) Correlation between the allometric exponent at TS and the number of spikelets per spike. (b) Correlation between the allometric exponent at GA and the maximum number of floret primordia. (c) Correlation between the allometric exponent at GA and the percentage of floret primordia that ended up setting grains (d) Correlation between the allometric exponent at GA and the number of grains per spikelet. (e) Correlation between the allometric exponent at YA and the harvest index at maturity.

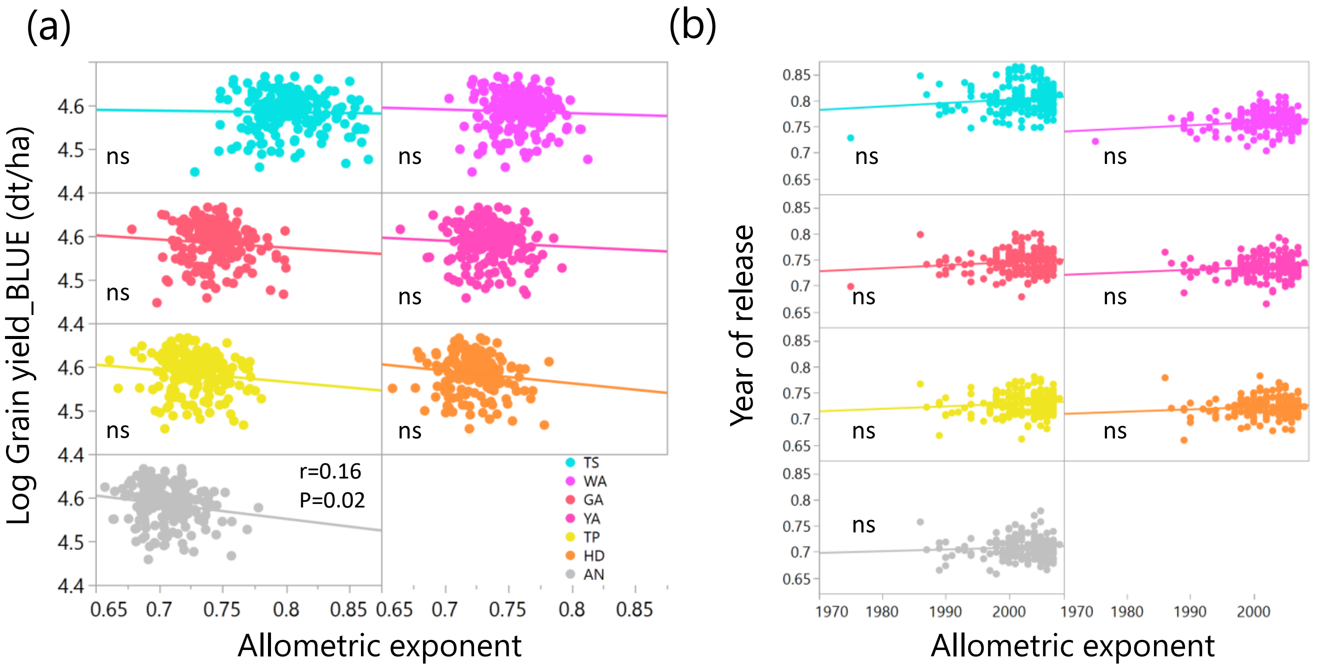


**Fig. S6** Growth allometry do not affect stable grain yield effects. (a) Relationships between the allometric exponent at different stages and the cultivar's grain yield per area, averaged across eight environments. (b) Relationships between the cultivar's year of release and the allometric exponent at different developmental stages.

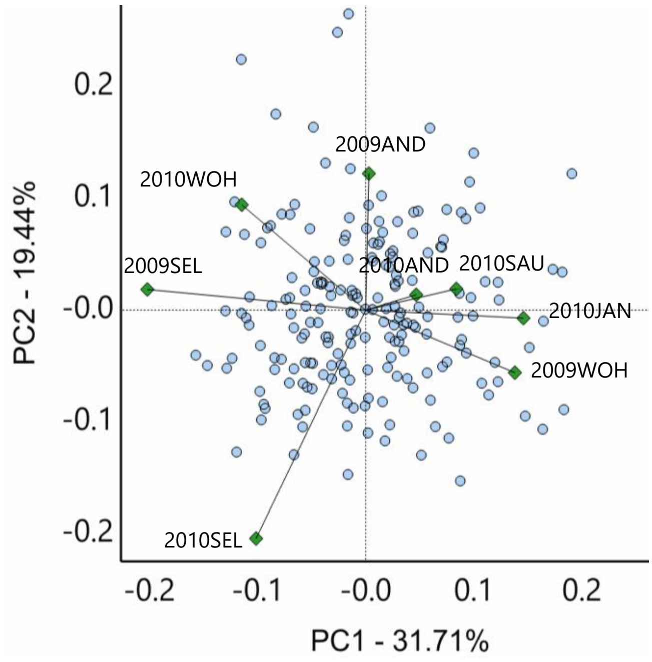


**Fig. S7** Visualization of G × E interactions for grain yield across eight environments in Germany and France. A biplot using symmetric scaling represents cultivars as blue circles and environments as green diamonds. Cultivars near the origin are less sensitive to environmental interactions, whereas those located farther from the origin exhibit specific adaptation.

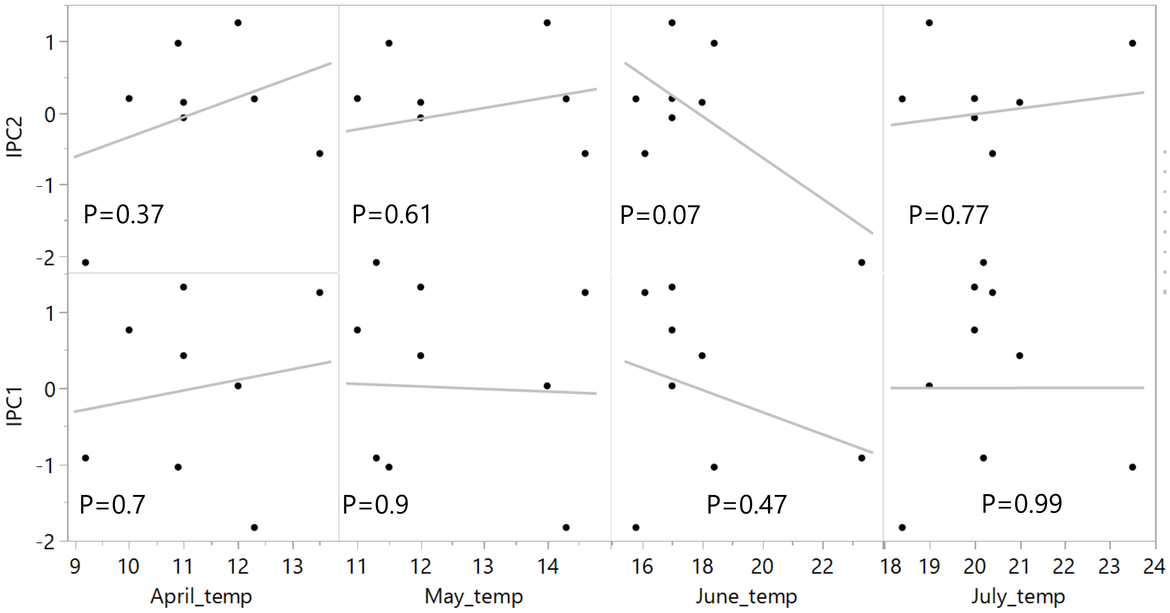


**Fig. S8** Correlations between the average temperature(Celsius degrees) in the eight field trials with the interaction principal components (IPC) extracted from the AMMI model.

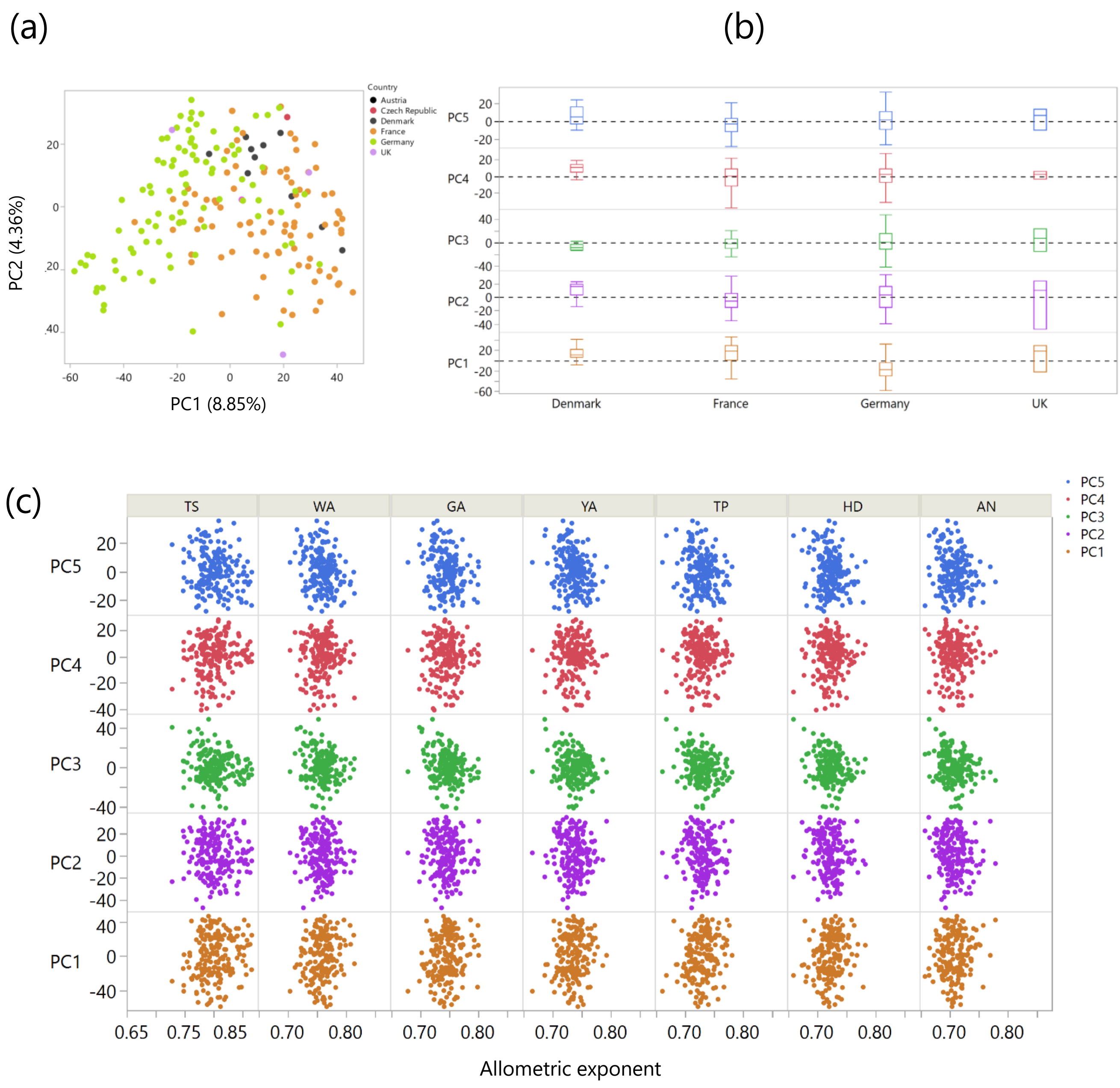


**Fig. S9** Population structure of the GABI wheat panel and its association with the country of origin and the scaling exponent. (a) Biplot of the population structure of the GABI wheat panel. (b) Association between the cultivar's country of registration and the population structure principal components. (c) Relationships between the scaling exponent at different stages and the population structure principal components.

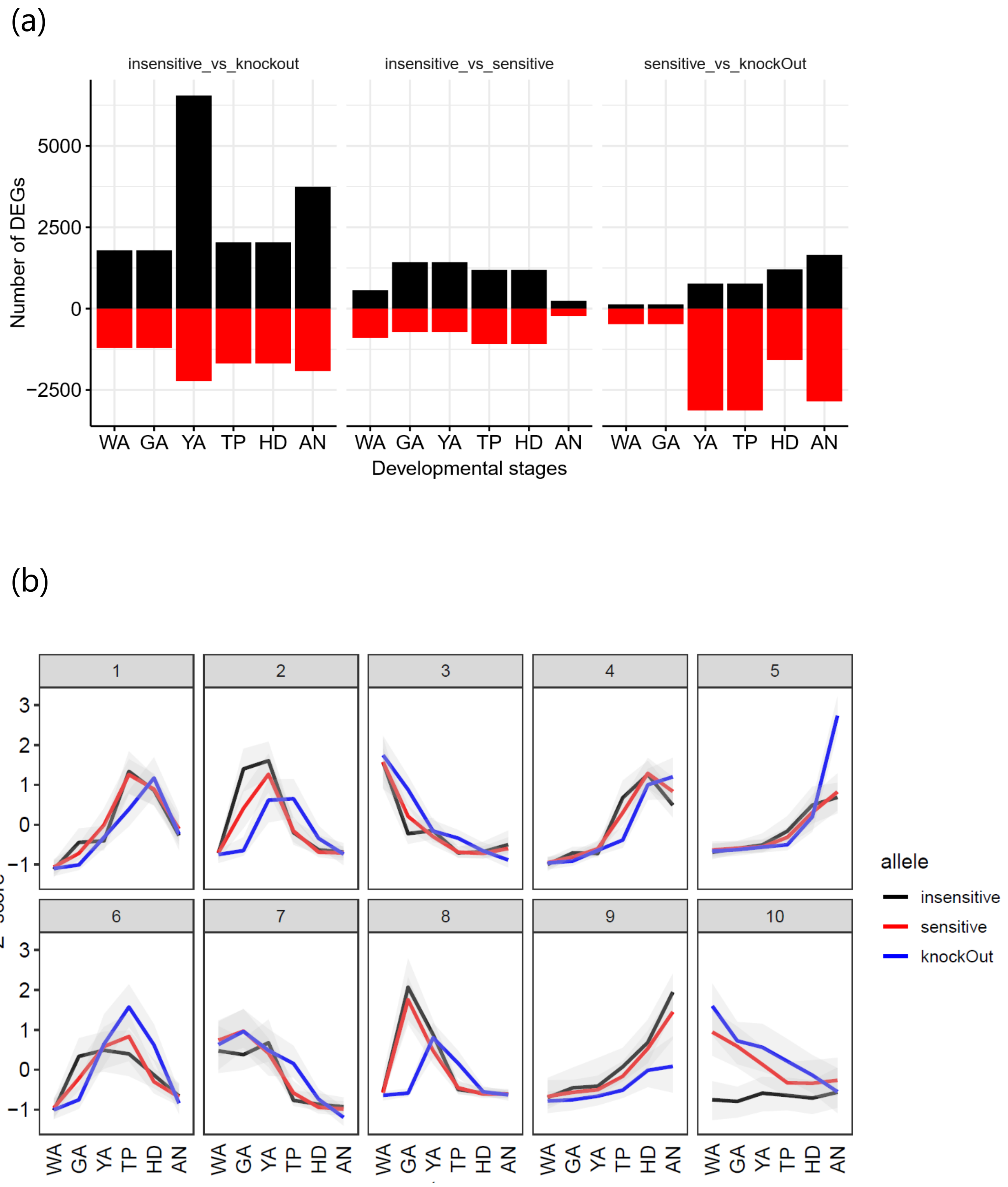


**Fig. S10** Gene expression in the spikelets of near-isogenic lines with differing *Ppd-1* alleles, observed from WA to AN. (a) The number of up regulated and down regulated genes between genotypes at different developmental stages. (b) Clusters of DEGs expression during spikelet growth.
